# A critical period terminates the differentiation of olfactory sensory neurons

**DOI:** 10.1101/2020.02.12.945428

**Authors:** Shadi Jafari, Mattias Alenius

**Affiliations:** Department of Clinical and Experimental Medicine, Linköping University, Linköping, Sweden; Department of Biology, New York University, New York, USA; Department of Molecular Biology, Umeå University, Umeå, SE 901 87, Sweden

## Abstract

Development generates a vast number of neuron types and classes. When and how neuronal differentiation end is poorly understood. Here, we show that *Drosophila* olfactory sensory neurons (OSNs) matures during a critical period and reveal that the differentiation termination mechanism is similar to the mammalian odorant receptor (OR) choice mechanism. We first showed that initiation of *Drosophila* OR expression required heterochromatin opening and a H3K9me3 demethylase, *Kdm4b*. Further genetic studies demonstrated that *Lsd1* and *su(var)3-9*, similar to mouse, were required to balance heterochromatin in order to stabilize OR expression. Expression analysis showed that *Lsd1, su(var)3-9* increased and *Kdm4b* decreased during the first two days after eclosion. We further showed that environment changes during the period, but not after, caused permanent transformed Lsd1, su(var)3-9 and Kdm4b expression and altered OR gene regulation. These results together suggest the last step in OSN terminal differentiation to be a gene regulatory critical period.

## Introduction

OSNs in most vertebrates and insects get specified to express a single odorant receptor (OR) from a large repertoire of OR genes in the genome and generate classes of OSNs that express a certain OR (Couto, Alenius, & Dickson, 2005; Fishilevich & Vosshall, 2005; Mombaerts et al., 1996; Ressler, Sullivan, & Buck, 1994). In *Drosophila*, it is assumed that the monogenic OR gene expression is strictly predetermined and non-stochastic (Barish & Volkan, 2015; Jafari et al., 2012; Ray, van Naters, & Carlson, 2008). There are several reasons for this assumption. For one, OR expression is stereotyped organised (Couto et al., 2005). *Drosophila* OSNs are also specified in a lineage dependent manner (Barish & Volkan, 2015) that is dictated by Notch and divide the post mitotic OSNs into two subgroups with defined projection patterns and expressed ORs (Endo, Aoki, Yoda, Kimura, & Hama, 2007). In addition to Notch and its effects, combinations of at least ten transcription factors (TFs) are required to express the different ORs (Jafari et al., 2012; Komiyama, Carlson, & Luo, 2004; Sim, Perry, Tharadra, Lipsick, & Ray, 2012; Tichy, Ray, & Carlson, 2008). However, while necessary, each TF combination is not sufficient to specify OR expression, as the defined TF combination is found elsewhere in the nervous system where the OR is not expressed (Jafari et al., 2012), suggesting that additional mechanisms restrict *Drosophila* OR expression.

Conversely, OR expression in vertebrates is considered to be less strict regulated and in part stochastic (Abdus-Saboor, Fleischmann, & Shykind, 2014; Monahan & Lomvardas, 2015). Cyclical changes in chromatin state from ‘active’ to ‘repressed’ and back again have been acknowledged as means for regulation of OR expression (Lyons et al., 2013; Magklara et al., 2011). In mouse and zebrafish, OR genes are embedded in constitutive heterochromatin in immature OSNs and in most non-olfactory tissues (Magklara et al., 2011). Constitutive heterochromatin is a condensed and transcriptionally inert chromatin conformation, marked by histone H3 lysine 9 trimethylation (H3K9me3) (Filion et al., 2010). Mathematical modelling of the vertebrate OR choice propose that the opening of constitutive heterochromatin around the OR gene initiate expression (Tan, Zong, & Xie, 2013) and requires a yet-to-be-identified H3K9me3 demethylase (Lyons et al., 2013). Next, Lsd1 remove the methyl groups from the OR locus marked with H3K9me2, which further open the chromatin and secure the OR expression initiation (Lyons et al., 2013). The folding or activity of the expressed OR induce several feedback loops that suppress other OR genes expression through down regulation of Lsd1 or upregulation of heterochromatin formation (Dalton, Lyons, & Lomvardas, 2013; Ferreira et al., 2014; Fleischmann, Abdus-Saboor, Sayed, & Shykind, 2013; Lyons et al., 2013).

*Drosophila* OR promoters are enriched with the H3K9me2 mark (Sim et al., 2012). Sim et al also show that su(var)3-9, which produce H3K9me3 and form constitutive heterochromatin, suppress spurious OR expression in *Drosophila*. Thermodynamic modeling based on detailed genetic studies of Or59b gene regulation put forward that cooperative interactions between transcription factors defy Su(var)3-9 suppression and promote *Drosophila* OR expression (Gonzalez, Jafari, Zenere, Alenius, & Altafini, 2019; Jafari & Alenius, 2015).

Here, we further address the role of heterochromatin in *Drosophila* OR regulation and show that similar to vertebrates, *su(var) 3-9* and *dLsd1* are a key factors in the commencement and maintenance of restricted OR expression. We reveal that the H3K9me3 demethylase, *Kdm4b*, is required to initiate *Drosophila* OR expression. We demonstrate that Kdm4b, dLsd1 and su(var)3-9 regulation is part of the closing mechanism of a gene regulatory critical period that marks the final step in the *Drosophila* OSN differentiation.

## Results

### *Kdm4b* initiate OR expression

Vertebrate OR expression is hypothesized to be initiated by an unknown factor that opens the constitutive heterochromatin at a single OR locus (Lyons et al., 2013; Tan et al., 2013). Our previous results demonstrate that whilst *Or59b* expression is stabilized by a constitutive heterochromatin configuration (Jafari & Alenius, 2015), computer modeling recently predicted that its opening is required for *Or59b* expression (Gonzalez et al., 2019). This event is initiated by H3K9me3 demethylation (Klose et al., 2006). In *Drosophila*, there are two H3K9me3 demethylases, *Kdm4a* (Kdm4B in vertebrates) and *Kdm4b* (Kdm4A, C, D, E in vertebrates) (Greer & Shi, 2012). UAS-IR (inverted repeats) for the two demethylases was expressed with *Peb-Gal4* in the OSNs and the knock down of the two demethylases showed that *Kdm4b* but not *Kdm4a* is required for *Or59b* expression (Figure 1A). The two Kdm4 enzymes have affinity for both H3K9 and H3K36 but Kdm4b is the major H3K9 demethylase (Tsurumi, Dutta, Shang, Yan, & Li, 2013), supporting that opening of heterochromatin initiate *Or59b* expression. To address if Kdm4b is required for continuous *Or59b* expression, we knocked down Kdm4b after OR initiation. As initiation of OR expression starts in the mid-pupal stages, we utilized *Orco-Gal4* which is expressed immediately before hatching (Larsson et al., 2004). *Orco-Gal4* driven Kdm4b knock down produced no changes in *Or59b* expression (Figure 1B), which indicate that Kdm4b is only required to initiate *Or59b* expression.

**Figure 1.**
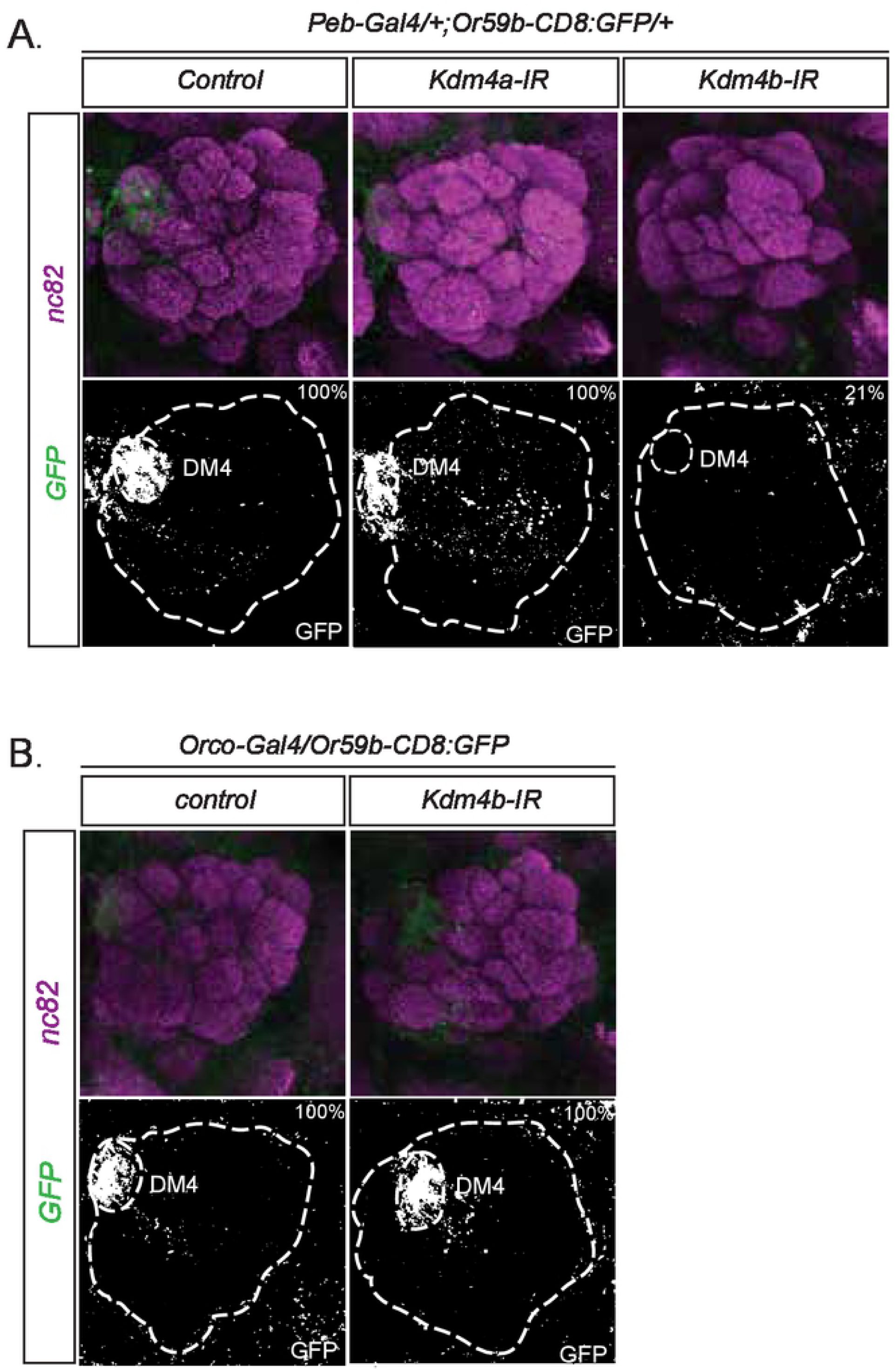
Kdm4b initiate Or59b expression. Whole-mount brain staining shows the expression of GFP (green) driven by the Or59b reporter. Synaptic neuropil regions are labeled with the presynaptic marker nc82 (magenta). Below each merged image, the GFP channel is shown. The marked region defines the antennal lobes and the DM4 glomerulus. (A) Loss of expression of the Or59b reporter is observed in knockdown of Kdm4b but not Kdm4a. Control flies were crossed to w1118. (B) Knock down of Kdm4b after the initiation of OR expression does not affect the expression of Or59b reporter.

### *dLsd1* and *Su(var)3-9* balance OR expression

Kdm4b converts H3K9me3 to H3K9me2 (Tsurumi et al., 2013). H3K9me2 in mouse OSNs is the substrate for Lsd1, which catalyse further demethylation to H3K9 and induce stable OR expression (Lyons et al., 2013). To investigate if dLsd1 is also required for *Drosophila* OR expression, we knocked down *dLsd1* in OSNs with *Peb-Gal4. Or59b* expression was lost in the knock down flies (Figure 2A). Analysis of two additional ORs showed that whilst *Or92a* expression was completely abolished in knock down antennae (Figure 2B, VA2), no change was observed for *Or47b* expression (Figure 2B, Va1v), demonstrating that dLsd1 is required for the expression of some but not all ORs.

**Figure 2.**
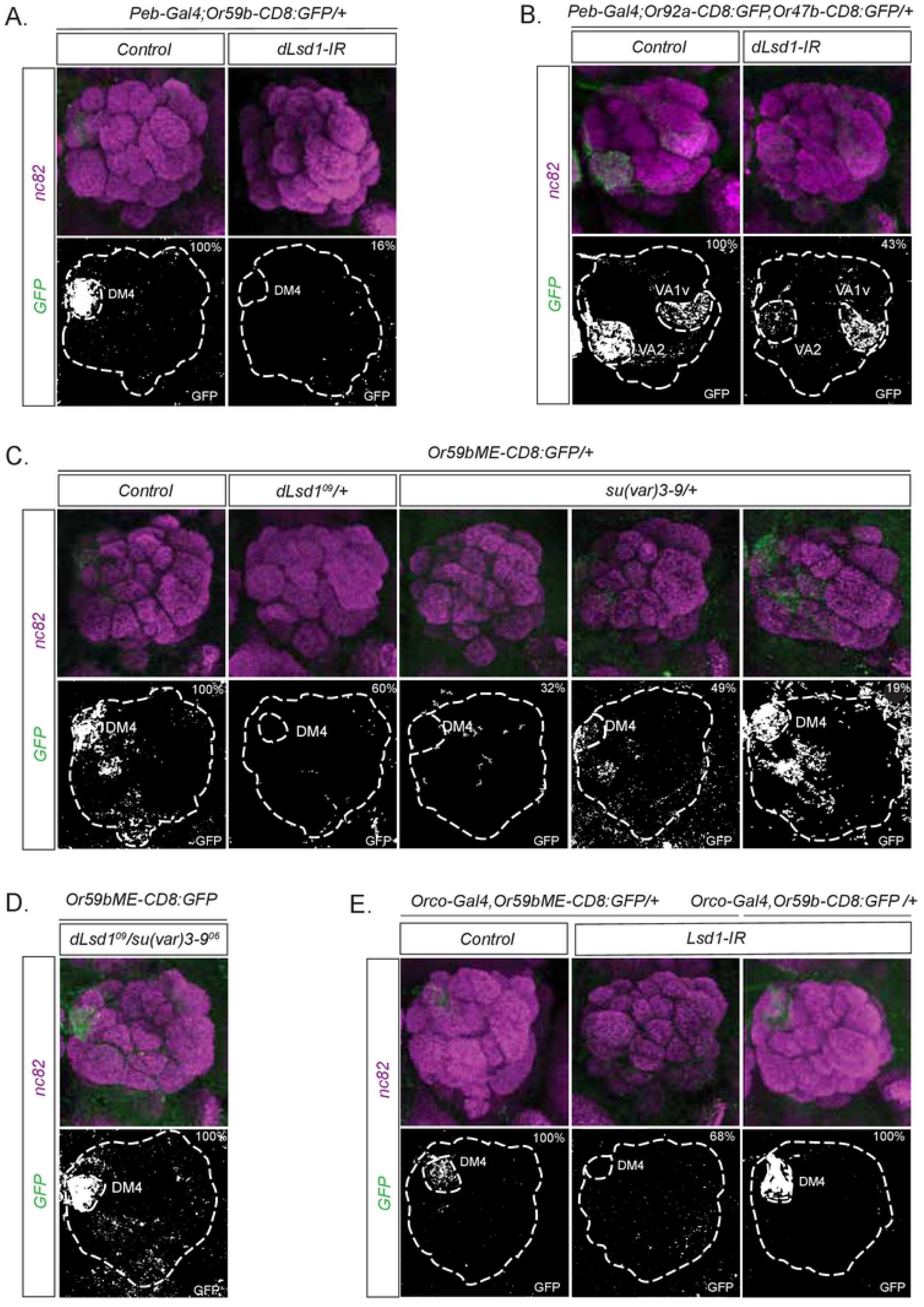
dLsd1 stabilize OR expression. Whole-mount brain staining shows the expression of GFP (green) driven by the OR reporter(s). Synaptic neuropil regions are labeled with the presynaptic marker nc82 (magenta). Below each merged image, the GFP channel is shown. AL and labeled glomeruli a are marked. (A) Or59b reporter expression is abolished in dLsd1 knockdown flies. Control flies were crossed to w^1118^. (B) Or92a and Or47b reporter GFP expression label the VA2 and VA1v glomeruli. Note the loss of Or92a reporter GFP expression in VA2 but not by the Or47b reporter (VA1v) in dLsd1 knock down flies. (C) GFP expression (green) driven by an Or59b minimal enhancer in su(var)3-9^06^ and dLsd1^09^ heterozygote flies. (D) The disturbed expression of the minimal enhancer in heterozygote flies is rescued double heterozygotes for su(var)3-9^06^ and dLsd1^09^. (E) Knock down of dLsd1 after the initiation of OR expression does not affect the expression of Or59b reporter but hampers the expression of the Or59b minimal enhancer.

During *Drosophila* development, dLsd1 promotes closing of heterochromatin, and not its opening (Di Stefano, Ji, Moon, Herr, & Dyson, 2007; Rudolph et al., 2007). In both *Drosophila* and mouse, heterochromatin formation is regulated by *Su(var)3-9*, which catalyses the methylation of H3K9me2 to H3K9me3 (Nakayama, Rice, Strahl, Allis, & Grewal, 2001; Rea et al., 2000; Schotta et al., 2002). To address if dLsd1 counteracts Su(var)3-9 activity during *Or59b* gene regulation, we reduced the gene copy number of either *dLsd1* or *Su(var)3-9*, or both. To sensitize the analysis we used a *Or59b minimal enhancer* reporter (*Or59bME*) that lack the cooperative transcription factor interactions required to defy heterochromatin formation (Jafari & Alenius, 2015). *Or59bME* reporter is expressed in the Or59b OSN class that project to the DM4 glomerulus but the reporter expression is sensitive to changes in chromatin state (Figure 2C (Jafari & Alenius, 2015). Consistently, in both *dLsd1* and *Su(var)3-9* heterozygote flies *Or59bME* reporter expression was lost (Figure 2C). Interestingly, combining the two heterozygotes rescued the single class expression of the reporter (Figure 2D). That restoring the copy number of the two genes complete rescued *Or59bME* regulation supports a model where dLsd1 opposes Su(var) 3-9 and promotes open heterochromatin in *Drosophila* OSNs. The results further demonstrate that strict coordination and balance of dLsd1 and Su(var)3-9 levels or function is required to support *Or59b* expression. Knock down after OR initiation of *dLsd1* resulted in loss of *Or59bME* expression, but not that of *Or59b* itself (Figure 2E), supporting that *dLsd1* continuously counter act heterochromatin formation and that transcription factor cooperativity reduce its requirement in the adult state. Interestingly, Lhx2, one of few transcription factors known to regulate vertebrate OR expression (Kolterud, Alenius, Carlsson, & Bohm, 2004), also requires cooperativity to counteract local H3K9me3 and heterochromatin formation in the OSNs (Monahan et al., 2017).

### HP1b and dLbr is required for Or59b expression

Heterochromatin on a local level represses expression and on a global level function together with HP1 proteins as attachment sites for the inner nuclear membrane anchor proteins that compartmentalize the nucleus into hetero- and eu-chromatin domains (van Steensel & Belmont, 2017; Wong, Luperchio, & Reddy, 2014). Nuclear localization of the vertebrate OR loci require the lamin b receptor (Lbr1) (Clowney et al., 2012). Knock down of *Drosophila dLbr1* in OSNs by *Peb-Gal4* drastically reduced both *Or59b* and minimal enhancer reporter expression (Figure 3A, B). Lbr1 interacts with HP1 proteins (Polioudaki et al., 2001; Ye & Worman, 1996). Knock down of HP1b but not HP1a, HP1c nor HP1d caused loss of *Or59b* expression (Figure 3B and Figure S1). Expression of the minimal enhancer reporter was also lost in both dLbr1 and HP1b knock down flies (Figure 3A). The *Or59b* reporter and the minimal enhancer reporter are inserted at different locations in the genome, which suggest thatthe OR reporters instructed the HP1 and dLbr1 function independent of locus location. Dynamic Kdm4b, dLsd1 and Su(var)3-9 expression modulate Or59b expression.

**Figure 3.**
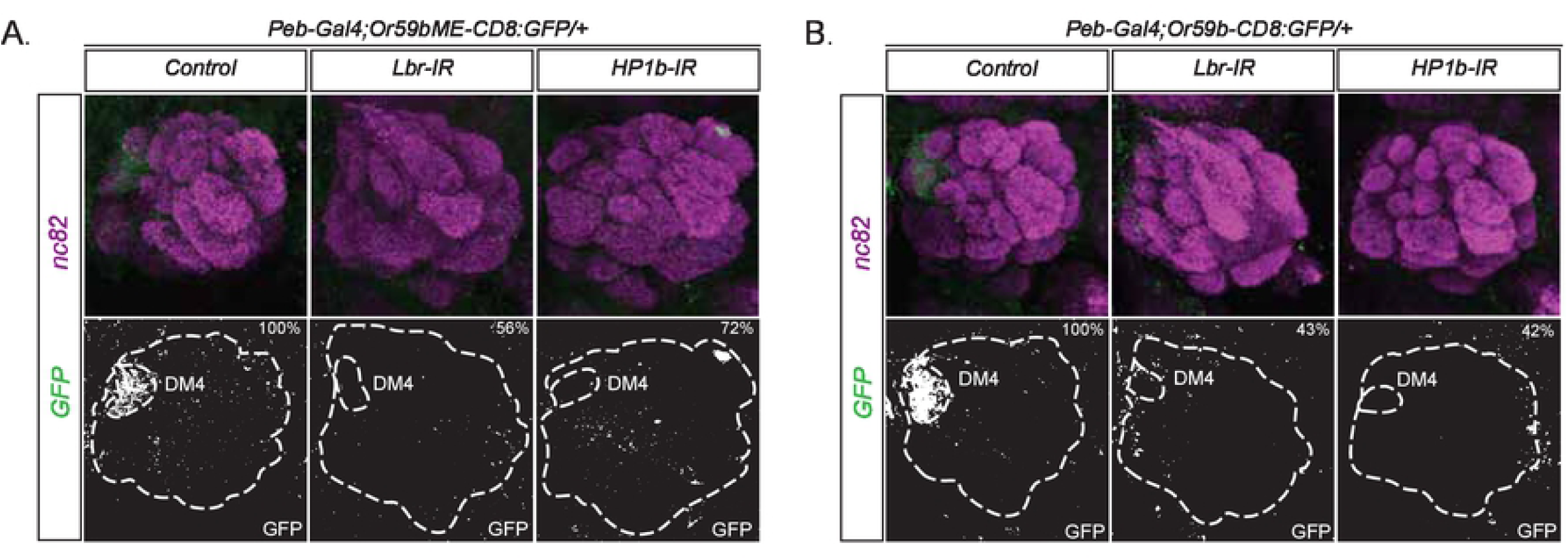
Lbr and HP1b defines the state of adult OR expression. Whole-mount brain staining shows the expression of GFP (green). Synaptic neuropil regions are labeled with the presynaptic marker nc82 (magenta). The antennal lobe and the DM4 glomerulus are out lined in the GFP channel. GFP expression (green) driven by the Or59bME reporter (A) and the Or59b reporter (B) is abolished in Lbr and HP1b knockdown flies. Control flies were crossed to w^1118^.

In immature flies, *Or59b* expression varied in level between OSNs and during the first days matured to the stable uniform expression level observed in adults (Figure 4A). Quantitative PCR (qPCR) experiments showed that *dLsd1* and *Su(var)3-9* expression increased in the antennae during the first days post eclosion (DPE), peaking in three-day old flies (Figure 4B), correlating with the maturation in OR expression. A more detailed analysis showed that the main increase in expression occurred during the first hours post eclosion (Figure 4C), which suggests that early exposure to odors and other environmental cues can modulate *dLsd1* and *Su(var)3-9* expression. The observation that *Kdm4b* expression decreased during the first days and remained low in adult flies (Figure 4D) further corroborates the idea that OR initiation and expression permissiveness decrease during a restricted time in early Drosophila life.

**Figure 4.**
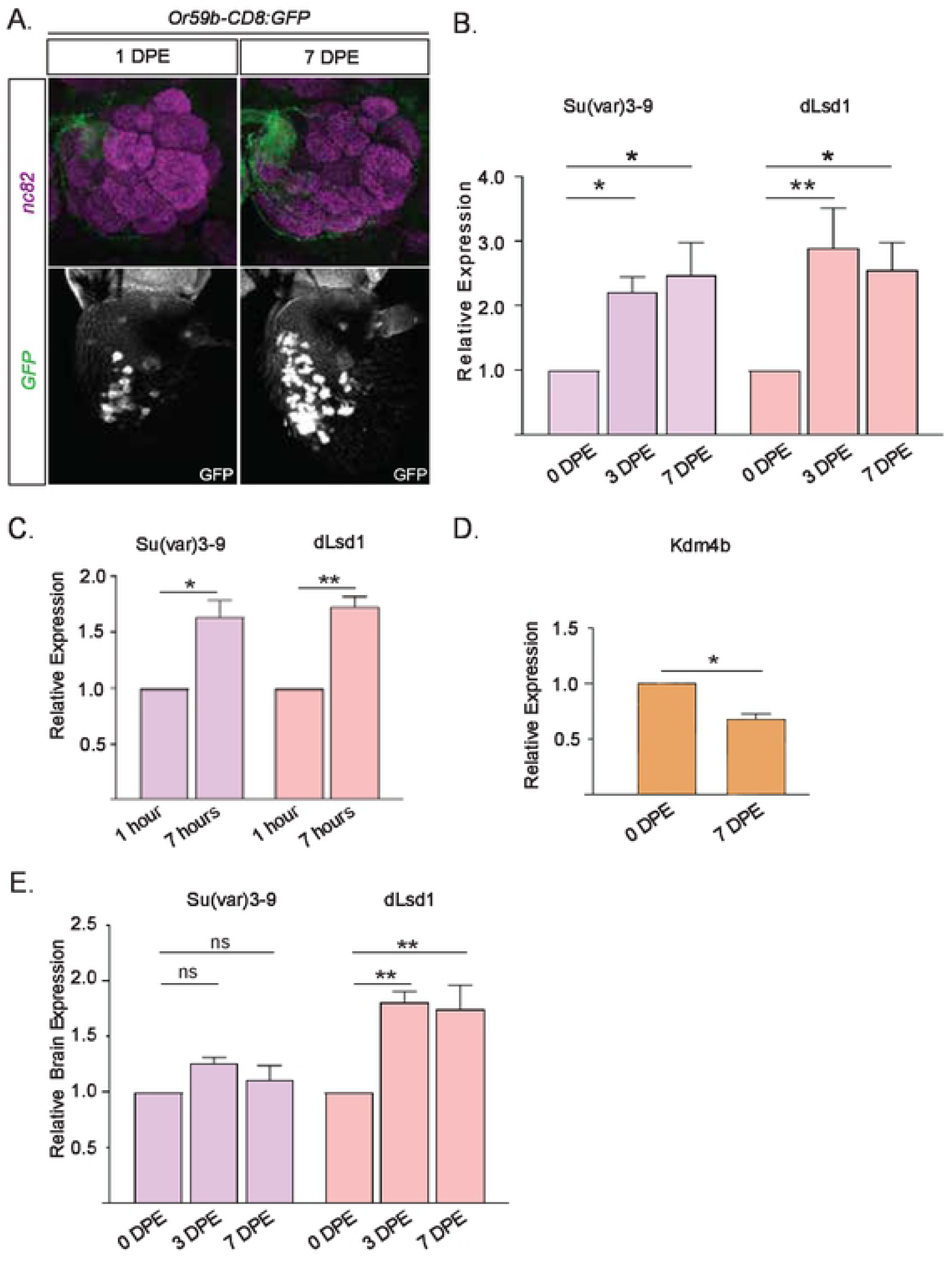
Or59b expression matures and dLsd1, Su(var)3-9and Kdm4b show dynamic expression in early-life Drosophila. (A) Whole-mount brain and antenna staining shows the expression of GFP (green) driven by the *Or59b* reporter. Synaptic neuropil regions are labeled with the presynaptic marker nc82 (magenta). Below each merged image, the GFP expression in the antenna is shown as the white channel. DPE, days post eclosion. (B) qPCR analysis of Su(var)3-9and dLsd1 expression in antenna at 0, 3 and 7 DPE (* p < 0.05; ** p < 0.01; *** p < 0.001; error bars represent SEM). (C) qPCR measurement of the Su(var)3-9 and dLsd1mRNA levels in antenna, 1 hour and 7 hours after eclosion. Note that the expression levels increase to almost twice in the first seven hours of adult fly’s life. (D) The mRNA levels of the Kdm4b in the antenna was measured by qPCR and compared between DPE 0 and 7. Note that unlike Su(var)3-9 and dLsd1, Kdm4b mRNA levels decrease post eclosion. (E) qPCR analysis of Su(var)3-9 and dLsd1 expression level in the brain at DPE 0, 3 and 7. Note that only dLsd1 increase in the brain post eclosion.

To address if *dLsd1* and/or *Su(var)3-9* expression also change in the *Drosophila* central nervous system, we performed qPCR on cDNA from brains of newly eclosed, three and seven-day old *Drosophila*. Interestingly, during the first three days *dLsd1* expression increased to the adult level (Figure 4E), showing that a similar general shift in *dLsd1* expression occurs in the brain. Surprisingly, *Su(var)3-9* expression was not altered (Figure 4E), indicating that the mechanism and change in gene regulation in the brain is different to the OSNs.

### *dLsd1* and *su(var)3-9* expression changes are a part of an OSN gene regulatory critical period

The well-defined and restricted temporal window of the refinement in OR expression (figure 4A) and change of *su(var)3-9, dLsd1* and *Kdm4b* expression resemble a critical period. If the observed changes is part of a critical period, the system should be sensitized to changes in the environment and neuronal activity during this time (Hensch, 2004). To address if the observed changes in *dLsd1* and *Su(var)3-9* expression is sensitive to environmental cues, we analysed changes to their levels in flies shifted from ambient to low temperature at different time points (Figure 5A). Flies subjected to a temperature shift at eclosion (1 DPE) showed a twofold reduction of *dLsd1* and *Su(var)3-9* expression (Figure 5B). Interestingly, a similar shift in adult flies (7 DPE) did not affect *Su(var)3-9* expression but *dLsd1* expression dropped to the level found at eclosion (Figure 5B). Thus, thermal stress differentially regulates *dLsd1* and *Su(var)3-9* expression in immature and adult flies, supporting the notion that modulation of *dLsd1* and *Su(var)3-9* expression takes place during a critical period of OSN gene regulation.

**Figure 5.**
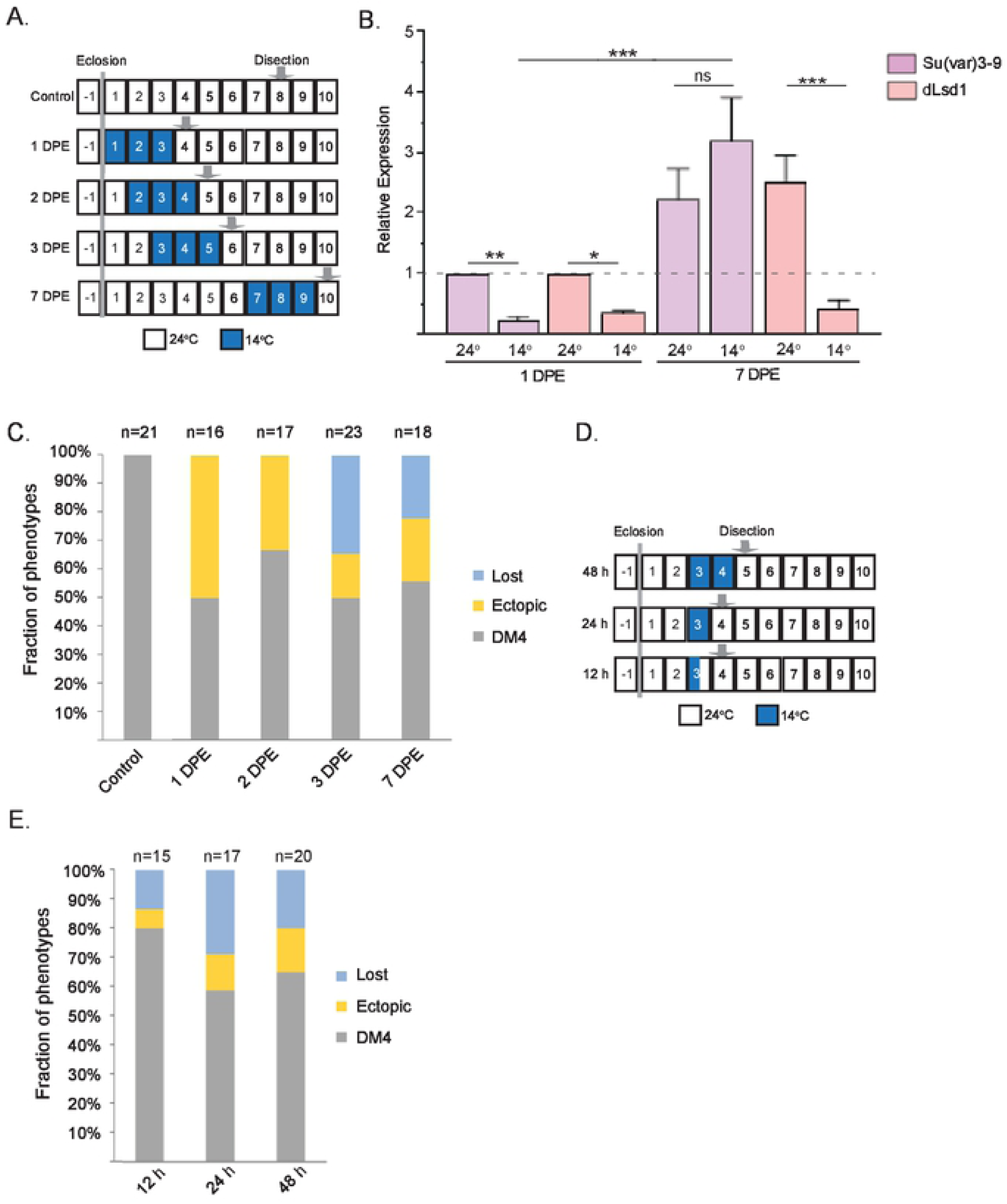
Environment changes during and after the critical period have contrasting effects on OR gene regulation, dLsd1 and Su(var)3-9expression. (A) Diagram decapitating the time course of experiments and low temperature shifts. Each box represents 24 hours. Low temperature exposure time is presented as dark blue boxes. (B) qPCR analysis of su(var)3-9and dLsd1 expression in antenna after low temperature shifts at DPE 1 or 7. (* p < 0.05; ** p < 0.01; *** p < 0.001; error bars represent SEM). (C) The percentage fractions of *Or59bME-GFP* brains that show control, ectopic or loss of expression after exposure to low temperature at DPE 1,2,3 or 7. (D) Diagram describing the restricted low temperature shifts. (E) The percentage fractions of 59bME reporter brains that showed control (DM4), ectopic or loss of GFP expression after the low temperature shifts denoted in D.

To visualize the effect of these events on OR gene regulation, we returned to the *Or59bME* reporter. At ambient temperature the *Or59bME* reporter was expressed in the Or59b OSN class that projects to the DM4 glomerulus (Figure 5C). *Or59bME* reporter flies shifted to low temperature immediately after eclosion showed a marked ectopic reporter expression with several GFP positive glomeruli (Figure 5C), which is consistent with the observed lowered *Su(var)3-9* and *dLsd1* expression (Figure 5B). Temperature shifts in adult flies produced mainly loss of expression (Figure 5C), which also tallies in with the observed imbalance in *Su(var)3-9* and *dLsd1* expression (figure 5B). Shorter shifts to low temperatures showed that the switch in reporter expression regulation occurred between day 2 and day 3 post eclosion (Figure 5D, E). Altogether, these results showed that there is a critical period in OSN gene regulation and that it is limited to the first two days’ post eclosion.

### Temperature stress during the critical period induce permanent changes in OSN gene expression

According to the defining criteria of a critical period, the state of the system at the end of the period should be permanent and irreversible (Hensch, 2004). To investigate if this is the case for *Or59bME* regulation, newly eclosed reporter flies were shifted to low temperature for three days and then returned to the ambient temperature for seven days. Consistent with a permanent change, the ectopic *Or59bME* reporter expression pattern persisted the seven-day recovery period (Figure 6A). Even after a prolonged 18-day recovery period the *Or59bME* expression pattern was comparable with that observed directly after the intervention (Figure 6A), demonstrating that the change in gene regulation during the first days was irreversible.

**Figure 6.**
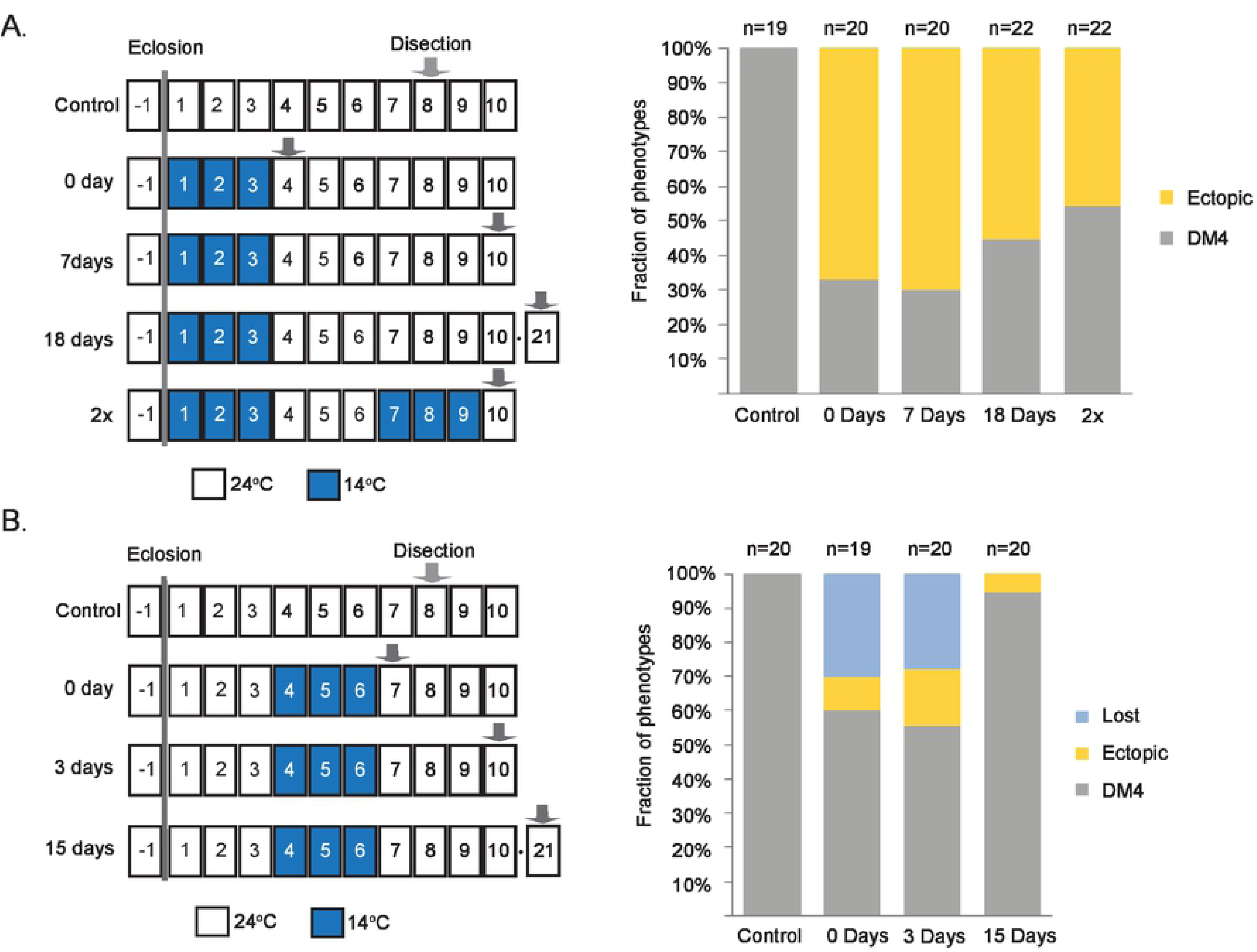
Environmental stress during the critical period induce permanent changes in OSN gene regulation. (A) The percentage fractions of the brains showing control or ectopic expression of *Or59bME-GFP* in the brain after exposure to low temperature at the day of eclosion for a period of three days. Note that neither a recovery of 18 days at 24°C nor a second exposure to low temperature from day 7 after eclosion affect the expression phenotype of the minimal enhancer. (B) The percentage fractions of the brains showing control, ectopic or loss of Or59bME reporter GFP expression in the brain after exposure to low temperature four days after eclosion for a period of three days. Note that a period of 14 days at 24°C recovers the phenotype to single OSN class expression.

If the critical period induces a transcriptional memory, the loss of *Or59bME* reporter expression observed after an adult temperature shift should recover to the expression pattern at the end of the critical period. Consistently, if 7 day old *Or59bME* reporter flies were shifted to low temperatures for three days, the loss of expression recovered to single OSN class expression (Figure 6B), demonstrating that temperature shifts after the critical period have only transient effects on the *Or59bME* reporter expression. To test if the critical period in fact produces a transcriptional memory and an irreversible permanent state, we challenged the ectopic expression of an early shift by a second adult shift to the same low temperature conditions. Strikingly, the ectopic expression pattern of the ME reporter was comparable between flies subjected to one or two shifts (Figure 6A, B). This demonstrated that the temperature during the critical period dictated *Or59bME* expression pattern in adult, thus producing a lasting transcriptional memory.

## Discussion

Here, we demonstrate that OSNs transition from a deterministic differentiation that produce variable OR expression levels to the mature stable high OR expression level observed in adult flies during a short window early in *Drosophila* life. This transition has the hallmarks of a critical period (Hensch, 2004) with a sharply-defined temporal window, sensitivity to external modulation, in this case environmental stress and generation of a permanent adult state, a terminal differentiated neuron.

Our findings further demonstrate that OSN differentiation during the critical period utilizes a mechanism similar to vertebrate OR choice. We first show that, as predicted for vertebrate OR expression (Tan et al., 2013), a H3K9me3 demethylase, Kdm4b initiates *Drosophila* OR expression. We further show that similar to the role in vertebrate OR expression, dLsd1 establish ORs expression in *Drosophila.* Our results and several vertebrate OR choice studies (Coleman, Lin, & Schwob, 2017; Lyons et al., 2013; Vyas, Meredith, & Lane, 2017) show that in the OSNs Lsd1 act as an activator that opposes Su(var)3-9 and constitutive heterochromatin formation. In most other *Drosophila* and vertebrate cells, Lsd1 function as a repressor that demethylates H3K4me1/2 and induces heterochromatin formation (Di Stefano et al., 2007; Rudolph et al., 2007), indicating that the distinct Lsd1 function in OSNs is a conserved feature.

The criteria for a critical period (Hensch, 2004) suggest that neuronal activity and competition is also part of the Drosophila OR regulation. Interestingly, there is evidence for competition between ORs in *Drosophila* (Shaw, Johnson, Anderson, de Bruyne, & Warr, 2019). Or22a and Or22b are two ORs expressed in a single class (Couto et al., 2005). In the wild *Drosophila melanogaster* express either Or22a or Or22b but not both, suggesting that there is an active suppression mechanism that maintain monogenic expression. In immature vertebrate OSNs, there is a low frequency of OR co-expression and biallelic expression (Hanchate et al., 2015; Shykind et al., 2004). To secure monogenic OR expression several feedback loops later suppress the spurious OR expression and restrict expression to the first or perhaps most expressed OR (Abdus-Saboor et al., 2014). Refinement feedback mechanisms build on that folding (Dalton et al., 2013; Lyons et al., 2013) or activity of the expressed OR (Ferreira et al., 2014; Fleischmann et al., 2013) suppress other OR genes expression through down regulation of Lsd1 or upregulation of heterochromatin formation. Thus, it will be of great interest to investigate if there are feedback mechanisms that link *Drosophila* OR expression or OSN activity with *Su(var)3-9* and *dLsd1* expression and regulate the critical period.

After its initiation, OR expression regulation differs between *Drosophila* and mouse. The most dramatic difference is that mouse ORs are mono allelic expressed whereas *Drosophila* ORs are like most genes biallelic expressed. Our results show that continuous *Drosophila* OR expression requires both *dLsd1*and *Su(var)3-9* and that heterochromatin state is under constant regulation, which underscore the difference to vertebrates where Lsd1 is down regulated after the initiation of the mono allelic expression. Together with the discovered critical period the constant regulation of heterochromatin implies that there must be a transcription memory mechanism, which sustain continuous OR expression. Our results together with recent published results indicate that nuclear organisation can be part of such a memory mechanism. We show here that *HP1* and *dLbr* are required for OR expression. Lbr and HP1 interacts (Polioudaki et al., 2001; Ye & Worman, 1996) and recent ChIP experiments show that in *Drosophila* neurons actively transcribed gene regions have a HP1/H3K9me2 profile (Pindyurin et al., 2018). A model where HP1 attach H3K9me2 heterochromatin with LBR and like that stabilize the nuclear organization of the OSN and OR expression could thus explain the somewhat surprising requirement for heterochromatin in Or59b expression. The need to coordinate H3K9me2 and heterochromatin levels is further underscored by that stable OR expression require the H3K9me2 methylase G9a in both mouse and Drosophila (Alkhori, Ost, & Alenius, 2014; Lyons et al., 2014).

Our qPCR results reveal an increase in *dLsd1* expression during the first days in the brain. A change in H3K9me2 regulation can thus be part of a potential general critical period in the *Drosophila* brain. Interestingly, there is no change in *Su(var)3-9* expression, which suggest that in early-life Drosophila the contrast in gene regulation between central neurons and OSNs increase. Thus, one direct implication of the lack of *Su(var)3-9* change is that it might suppress OR expression in the central nervous system, which would be an elegant and simple solution to limit OR expression to the OSNs. Still, if there is a general critical period in terminal differentiation of Drosophila neurons and how *dLsd1* and *Su(var)3-9* expression is regulated in central and sensory neurons are interesting questions that remains to be addressed.

## Materials and Methods

### Drosophila stocks

Or59b promoter fusion and Or59b minimal enhancer constructs have previously been described (Couto et al., 2005; Jafari & Alenius, 2015). The *Pebbled-Gal4* (*Peb-Gal4*) was a kind gift from Liqun Luo (Stanford University, Stanford, CA, USA). The *su(var)3-9*^*06*^ and *Lsd1*^*09*^ mutants were a kind gift from Anita Öst (Linköping University, Linköping, Sweden). The following fly lines were provided by the Bloomington Drosophila Stock Center: *w*^*1118*^ (38690), *Orco-Gal4* (23909), *su(var)205*^*5*^ (12226) *dLsd1-IR* (36867; 32853, 33726), *dLbr-IR* (53269), *Hp1b-IR* (32401), *Hp1c-IR* (31339, 33962), *su(var)205 (Hp1a)-IR* (33400), *rhi (Hp1d)-IR* (34071, 35171) *Kdm4a-IR* (34629), *Kdm4b-IR* (35676, 57721).

Flies were raised and maintained on standard Drosophila culture medium at 24°C and collected at the day of eclosion and aged unless otherwise specified. Temperature shifted flies were transferred to new vials and maintained for 3 days at 14°C.

### Immunofluorescence

Immunofluorescence was performed according to previously described methods. The following primary antibodies were used: rabbit anti-GFP (1:2000, TP-401; Torrey Pines Biolabs) and mouse anti-nc82 (1:100; DSHB). Secondary antibodies were conjugated with Alexa Fluor 488 (1:500; Molecular Probes) and Goat anti-Mouse IgG (H+L) Cross-Adsorbed Secondary Antibody, Rhodamine Red-X (1:250), Thermo Fisher). Confocal microscopy images were collected on an LSM 700 (Zeiss) and analyzed using an LSM Image Browser. Adobe Photoshop CS4 (Adobe Systems) was used for image processing.

### qPCR

After freezing flies in liquid nitrogen antennae were obtained with a sieve. Total RNA from antennae was extracted with TRIzol reagent (Invitrogen) and the RNeasy kit (Qiagen). Quantitative PCR was conducted with an Applied Biosystems 7900HT real-time PCR system (Life Technologies) using the Power SYBR Green PCR master mix (Applied Biosystems, Life Technologies) and primer sets designed using Primer Express software v3.0.1 (Integrated DNA Technologies). Tubulin and Actin were used as an internal control for the experiments. To amplify cDNA products and not genomic DNA, primers were designed to join the end of one exon with the beginning of the next exon. Quantitative PCR for each primer set was performed on both control and experimental samples for 40 cycles. Following amplification, melt curve analysis and ethidium bromide agarose gel electrophoresis were performed to evaluate the PCR products. The relative quantification of the fold change in mRNA expression was calculated using the 2−ΔΔCT threshold cycle method.

## Acknowledgements

We thank, Anita Öst for flies; Najat Dzaki, Carlos Ribeiro and Staffan Bohm for discussions and comments on the manuscript. This work was supported by the Swedish Research foundation, grant (2016-05208)

